# Facilitation of neural responses to targets moving against optic flow

**DOI:** 10.1101/2020.06.28.172536

**Authors:** Sarah Nicholas, Karin Nordström

## Abstract

For the human observer, it can be difficult to follow the motion of small objects, especially when they move against background clutter. In contrast, insects efficiently do this, as evidenced by their ability to capture prey, pursue conspecifics, or defend territories, even in highly textured surrounds. We here recorded from target selective descending neurons (TSDNs) which likely subserve these impressive behaviors. To simulate the type of background optic flow that would be generated by the pursuer’s own movements through the world, we used the coherent motion of a perspective distorted sparse dot field. We show that hoverfly TSDN responses to target motion are suppressed when such background optic flow moves in the same direction as the target. Indeed, the neural responses are strongly attenuated against both translational sideslip as well as rotational yaw. More strikingly, we show that TSDNs are facilitated by background optic flow in the opposite direction to the target, if the target moves horizontally. Furthermore, we show that a small, frontal spatial window of background optic flow is enough to fully facilitate or attenuate TSDN responses to target motion. We argue that the TSDN response facilitation could be beneficial in modulating corrective turns during target pursuit.

**Significance statement:** Target detection in visual clutter is a difficult computational task that insects, with their poor resolution compound eyes and small brains, do successfully and with extremely short behavioral delays. We here show that the responses of target selective descending neurons are attenuated by background motion in the same direction as target motion, but facilitated by opposite direction background motion. This finding is important for understanding conspecific pursuit behavior, since these descending neurons likely control behavioral output. The facilitation that we describe would come into effect if the hoverfly is subjected to background motion in one direction, but the target it is pursuing moves in the opposite direction, and could therefore be used to modulate gaze stabilizing corrective turns.

## Introduction

The survival of many animals often depends on their ability to visually detect small moving objects or targets, as these could represent predators, prey, mates or territorial intruders (1). Efficient target detection is a computationally challenging task, which becomes even more difficult when done against visual clutter. Despite this, many insects successfully detect targets, followed by highly acrobatic pursuits, often in visually complex environments. For example, male *Eristalis tenax* hoverflies establish territories in foliage rich areas, on alert for intruders or potential mates and ready to engage in high-speed pursuit (2).

Initial target detection can be facilitated by behaviors that render the background stationary, thus making the target the only thing that moves. Many insects and vertebrates indeed visualize targets against the bright sky (3), or from a stationary stance, such as perching (4–6) or hovering (7, 8). However, as soon as the pursuer moves, its own movement creates self-generated widefield motion across the retina, often referred to as optic flow (9), or background motion (10). In addition to self-generated background optic flow, when a pursuer is subjected to involuntary deviations away from their flight path, for example by a gust of wind, this also generates background optic flow. Quickly correcting such unplanned course deviations is essential for successfully navigating through the world. For example, a flying insect being pushed sideways to the right by a gust of wind, will experience optic flow in the opposite direction. To correct for this, in response to leftward optic flow, the insect will use its wings to perform a corrective optomotor response to the left (11), and/or stabilize its gaze by moving its head (12). Recent evidence, suggests that efference copies suppress the visual neurons sensitive to optic flow, as the optomotor response could otherwise counteract voluntary turns (13).

During pursuit, the pursuer is thus subjected to visual motion that could originate from the independent motion of objects within the environment, from its own intentional movements, or from movement imposed by external forces, such as a gust of wind. Importantly, how the insect moves also affects the type of optic flow it will experience. For example, during translations distant features move slower than closer ones, whereas during rotations, all features move at the same angular velocity irrespective of distance from the observer (14). Many insects use the information such relative motion provides to estimate the distance to features in the surround. *Eristalis tenax* hoverflies, for example, make both small translational head movements when perched and larger side-to-side movements during flight (15, 16). Moreover, many flying insects, including *Eristalis’* show behavioral segregation between rotational and translational movements, through a saccadic flight and gaze strategy (16–18). How this may influence target detection is currently not known.

The ability of insects to successfully pursue targets in clutter (4, 19) is thus remarkable, and suggests a high level of optimization, making the underlying neural mechanisms interesting to study. Indeed, insects that efficiently pursue targets, including predatory dragonflies and robberflies, as well as territorial hoverflies, have higher-order neurons that are sharply tuned to the motion of small, dark targets, identified in the optic lobes (20, 21) and the descending nerve cord (22, 23). Target tuned neurons often have receptive fields (24–26) in the part of the compound eye that has the best optics (27, 28). Target selective descending neurons (TSDNs) project to the thoracic ganglia (25, 29) where wing and head movements are controlled (30), and electrically stimulating dragonfly TSDNs leads to wing movements (31). Taken together, this suggests that TSDNs subserve target pursuit. However, how TSDNs respond to targets moving against translational and rotational optic flow is unknown.

We quantified the responses of *Eristalis* TSDNs to targets moving against 6 types of optic flow, three translations and three rotations. The background optic flow consisted of the coherent motion of thousands of “targets”, so that the target could only be distinguished by its relative motion. We found that optic flow in the same direction as the target attenuated the TSDN response to the target, whether the optic flow moved in a rotational or translational manner. We found that perpendicular optic flow attenuated the TSDN target response, but to a lesser degree than syn-directional background motion. This suggests that the vector divergence between the target and the background is important. Most strikingly, we found that optic flow in the opposite direction to target motion increased the TSDN response, if the target moved horizontally. Such neural facilitation is a novel observation. We found that a small spatial extent of frontal motion was sufficient to elicit both TSDN attenuation and facilitation, whether spatially overlapping with target motion or not. As descending neurons control behavioral output (30, 31), the response attenuation and facilitation could play a role in modulating optomotor or gaze stabilizing corrective turns as needed during target pursuit.

## Results

### TSDNs respond to target motion but not to background optic flow

We simulated background optic flow using the coherent motion of a sparse dot field (24). By projecting the individual features in the sparse dot field onto a screen anterior of the hoverfly, their spatial location, and simulated z-depth, over time, provided the type of optic flow that would be generated by self-motion (24). In optic flow sensitive descending neurons, this stimulus elicits strong direction-selective responses, similar to the responses to widefield sinusoidal gratings (24), or large moving images (22) with naturalistic statistics (32). We confirmed that our optic flow stimulus did not generate a response in target selective descending neurons (TSDNs) in male *Eristalis tenax* hoverflies (Fig. S1). For example, translational sideslip to the left only generated a response in 18% of 222 repetitions across 12 TSDNs (Fig. S1A). Furthermore, when leftward sideslip did generate a TSDN response, this was much less than the response to a target traversing a white background (Fig. S1B). This lack of TSDN response confirms that the optic flow stimulus is qualitatively similar to other types of widefield background motion.

In contrast to the lack of response to the optic flow, TSDNs responded strongly to the motion of a small, dark target (22, 24) traversing a white background (compare spiking response during stimulation with the lack of activity before stimulation, Fig. 1, left, Movie 1, 2), as well as when the target traversed the stationary sparse dot field (Fig. 1, right, Movie 3, 4). Note, that since the sparse dot field consisted of hundreds of “targets”, there were no spatial characteristics separating the individual target from the optic flow when both were stationary (Fig. 1A, right). For quantification across neurons, we calculated the mean spike frequency for the duration of target motion, except for the first and last 40 ms of each 500 ms target trajectory (dotted box, Fig. 1B, C). We found a small, but significant response reduction to targets moving across a stationary dot field compared with a white background (Fig. 1D). As the response across neurons was variable (N=30, coefficient of variation 63% and 67%, respectively, Fig. 1D), we normalized the response from each neuron to its own mean response to a target moving across a white background.

**Figure 1.**
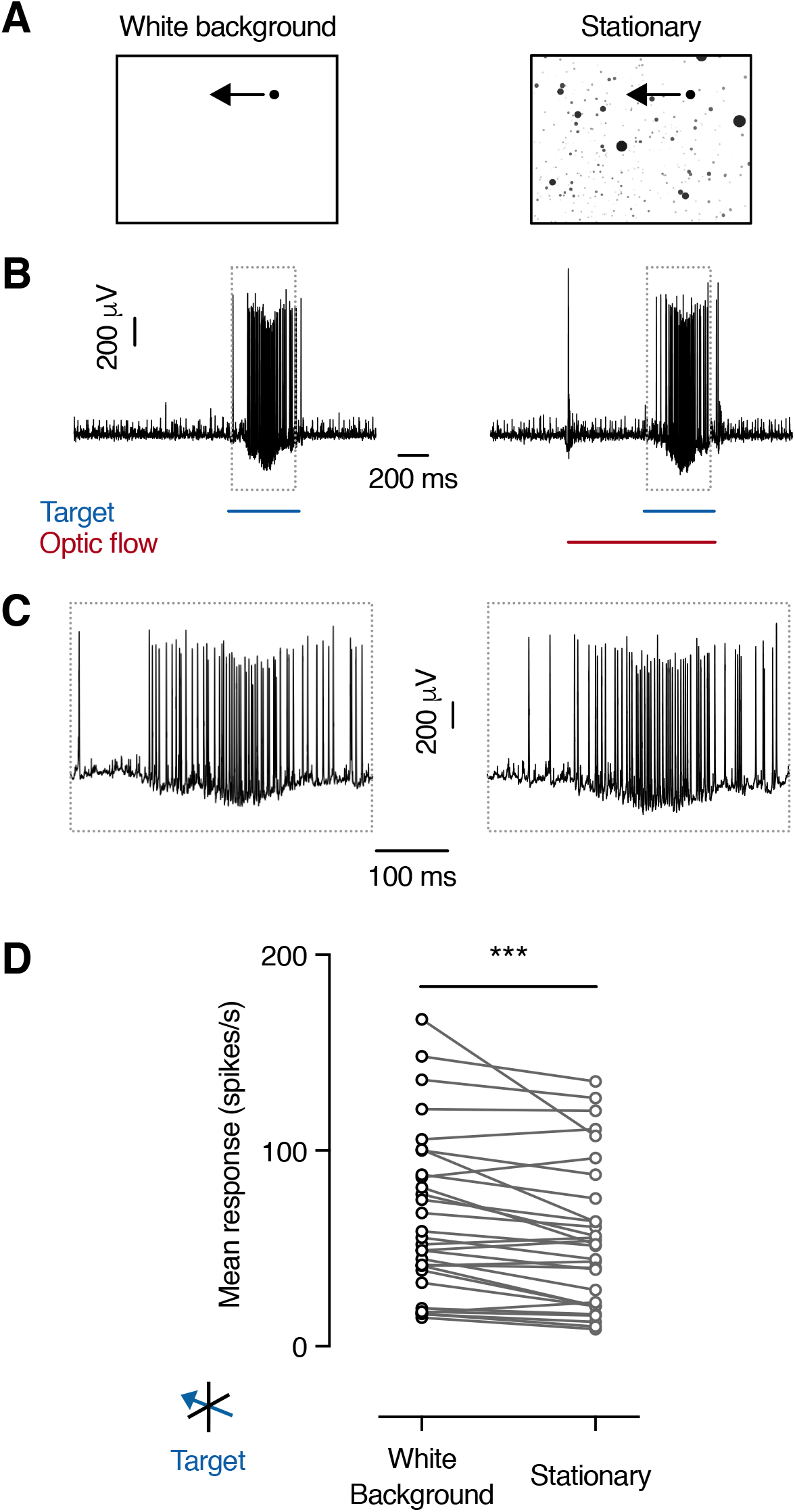
TSDN response to target motion. **A)** Pictograms of the round, black target, with a diameter of 3°, traversing a white background (left), or stationary optic flow (right), at 130°/s. **B)** Raw data trace from an extracellular TSDN recording. Timing of stimulus presentation indicated by colored bars (blue, target, and red, sparse dot field). **C)** Magnification of the raw data traces shown in panel B. **D)** The mean spiking response of different TSDNs was significantly reduced when the target moved across a stationary sparse dot field compared with a white background (N=30; ***P = 0.0005, two-tailed paired t-test).

### Background optic flow modulates the TSDN target response

To investigate the effect the background optic flow had on TSDN responses to target motion we simulated different types of translations (Fig. 2A) at 50 cm/s and rotations (Fig. 2B) at 50 deg/s. During translations distant features move slower than closer ones, whereas during rotations, all features move at the same angular velocity irrespective of distance from the observer (14). We first simulated translational sideslip and found that when the target moved horizontally across syn-directional sideslip, the TSDN response was strongly attenuated (Fig. 2C, D, “Left Sideslip”, Movie 5, 6), compared with control where the target moved over a stationary dot field (grey, Fig. 2C, D, Movie 3, 4). Similar effects of syn-directional background motion were previously seen when using panoramic images with naturalistic statistics (22).

**Figure 2.**
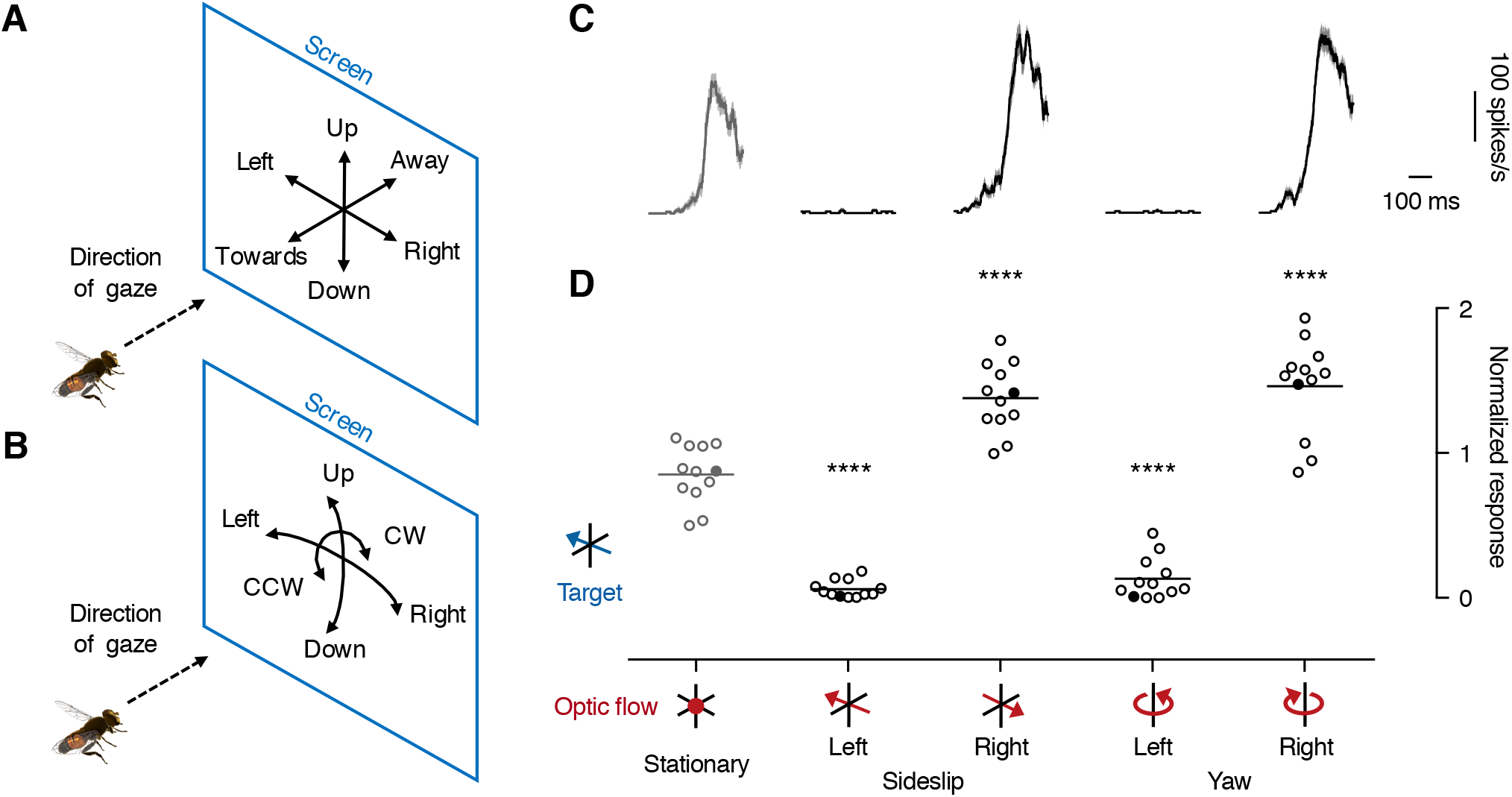
TSDN response to target motion is affected by horizontal background optic flow. **A)** A pictogram indicating the three different types of translations, as seen by the hoverfly. Sideslip was displayed to the left or the right, lift up or down, and thrust towards or away from the hoverfly, all at 50 cm/s. **B)** The three different types of rotations used. As these were simulated to rotate around the hoverfly’s position (24), the optic flow displayed on the screen can be described as yaw to the left or the right, pitch up or down, and clockwise or counter-clockwise roll, all at 50 deg/s. **C)** The TSDN response to a target traversing a stationary sparse dot field (left, grey), sideslip or yaw in either the same or the opposite direction to the target (red pictograms). All histograms from a single TSDN (mean ± sem, n=18) shown with 1 ms resolution after smoothing with a 20 ms square-wave filter. **D)** The mean response across TSDNs (N=12) to a target traversing stationary optic flow (left, grey), sideslip or yaw, in the same direction (Left”) or opposite direction (“Right”) to the target, after normalizing the data to each neuron’s own response to a target traversing a white background. The line shows the mean and the filled data points correspond to the neuron in panel C. In panel D significance is shown with **** for P < 0.0001 following a one-way ANOVA followed by Dunnett’s multiple comparison’s test done together with the data shown in Fig. 4.

To investigate the effect the relative motion of individual features within the optic flow had on TSDN responses to target motion, we next simulated rotational yaw at 50 deg/s. We found that yaw optic flow in the same direction as the target also strongly attenuated the TSDN response (Fig. 2C, D, “Left Yaw”). This suggests that rotational and translational optic flow have the same effect on TSDN responses to target motion, and that the relative motion of features within the optic flow is less important.

We next investigated the effect opposite-direction background optic flow had on TSDN responses to target motion. We found that when the target moved against counter-directional sideslip the TSDN response was strongly enhanced (Movie 7, 8, Fig. 2C, D, “Right Sideslip”, mean increase 71.1%). Similarly, yaw optic flow in the opposite direction to the target also facilitated the TSDN response, by 84.9% (Fig. 2C, D, “Right Yaw”). Such response facilitation was not seen when displaying targets over counter-directional background motion using panoramic images, in either TSDNs (22) or other target tuned neurons in the fly optic lobes (21, 26, 33).

### Frontal optic flow is required and sufficient

Our data show that yaw and sideslip have similar effects on TSDN responses to target motion (Fig. 2). As both sideslip and yaw contain substantial local motion in the frontal visual field (34), we next asked if frontal optic flow is required. We investigated this by limiting the spatial extent of the sideslip to either cover the ipsilateral, dorsal, ventral or contralateral position on the screen (Fig. 3A). Note that only the dorsal position covers the TSDN receptive field (Fig. 3A). In the other three positions, the sideslip optic flow was spatially separated from the receptive field (Fig. 3A), and thus the target trajectory.

**Figure 3.**
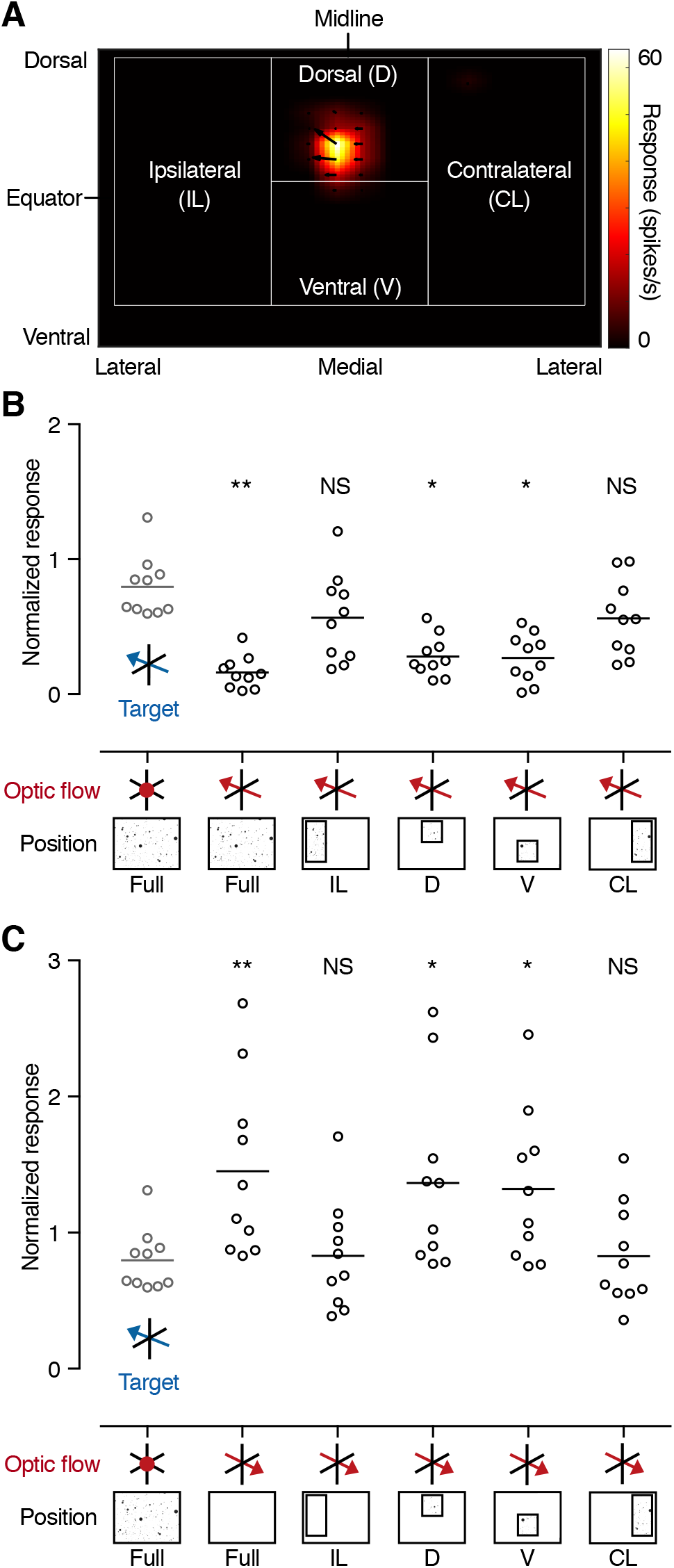
Frontal optic flow is required and sufficient. **A)** A pictogram of the separation of the background optic flow into four distinct positions: ipsilateral (IL), dorsal (D), ventral (V) and contralateral (CL). The color coding shows the receptive field of an example TSDN, and the arrows its local motion sensitivity. **B)** TSDN responses to targets moving across syn-directional sideslip are significantly inhibited compared to stationary control (grey) if the sideslip covers the full, dorsal or ventral screen. **C)** TSDN responses to targets moving across opposite direction sideslip are significantly facilitated compared to stationary control (grey, same data as in panel B) if the sideslip covers the full, dorsal or ventral visual field. Panels B and C show data from the same neurons (N=10, one-way ANOVA followed by Dunnett’s multiple comparisons test, with *P<0.05, **P<0.01).

We found that when sideslip moved in the same direction as the target, the TSDN response was attenuated if the optic flow covered the full, dorsal or ventral position on the screen (Fig. 3B), compared with the stationary control (grey, Fig. 3B). However, when the optic flow only covered the ipsilateral or contralateral positions, there was no difference compared with control (Fig. 3B), suggesting that frontal optic flow is both required and sufficient. Our results are consistent with previous work using moving images with naturalistic statistics, which did not have to spatially overlap with the target trajectory, nor have a large spatial extent, to attenuate TSDN responses (22).

We next found that when sideslip moved in the opposite direction to the target, the TSDN response to target motion was facilitated if the optic flow covered the full, dorsal or ventral positions of the screen (Fig. 3C). When the sideslip was limited to the ipsilateral or contralateral positions, there was no TSDN response facilitation (Fig. 3C). This suggests that the optic flow does not have to spatially overlap with the target trajectory. However, there has to be frontal, opposite direction optic flow for facilitation to take place.

In summary, our results show that a small spatial window of optic flow in either the dorsal or ventral visual field is enough to strongly attenuate (Fig. 3B) or facilitate (Fig. 3C) the TSDN response to target motion.

### Vector divergence between target and background optic flow affects TSDNs response

Our data above show that syn-directional optic flow strongly attenuates TSDN responses to target motion, whereas counter-directional optic flow facilitates the response (Fig. 2, 3). This suggests that the level of vector divergence between the target and the background influences the TSDN responses, so that maximum attenuation is generated at minimum vector divergence, whereas maximum facilitation is generated at maximum divergence. To explore this in further detail, we quantified the effect other types of optic flow had on TSDN responses to target motion.

We found that the TSDN target response was suppressed by lift, which moves perpendicular to the target (mean 44.0% response for downwards lift, “Lift Down”; mean 34.3% for upwards lift, “Lift Up”, Fig. 4). When the target was displayed against pitch, which also provides perpendicular motion in the frontal visual field, the response was suppressed to similar levels (57.1%, “Pitch Down”; 54.9%, “Pitch Up”, Fig. 4). The TSDN response to target motion was also attenuated when the target was displayed against thrust, which provides perpendicular motion along the animal’s anterio-posterior axis (44.3%, “Thrust Towards”; 58.4%, “Thrust Away”, Fig. 4). Thus, perpendicular background optic flow attenuates the TSDN response (Fig. 4), but not as much as syn-directional motion (Fig. 2).

**Figure 4.**
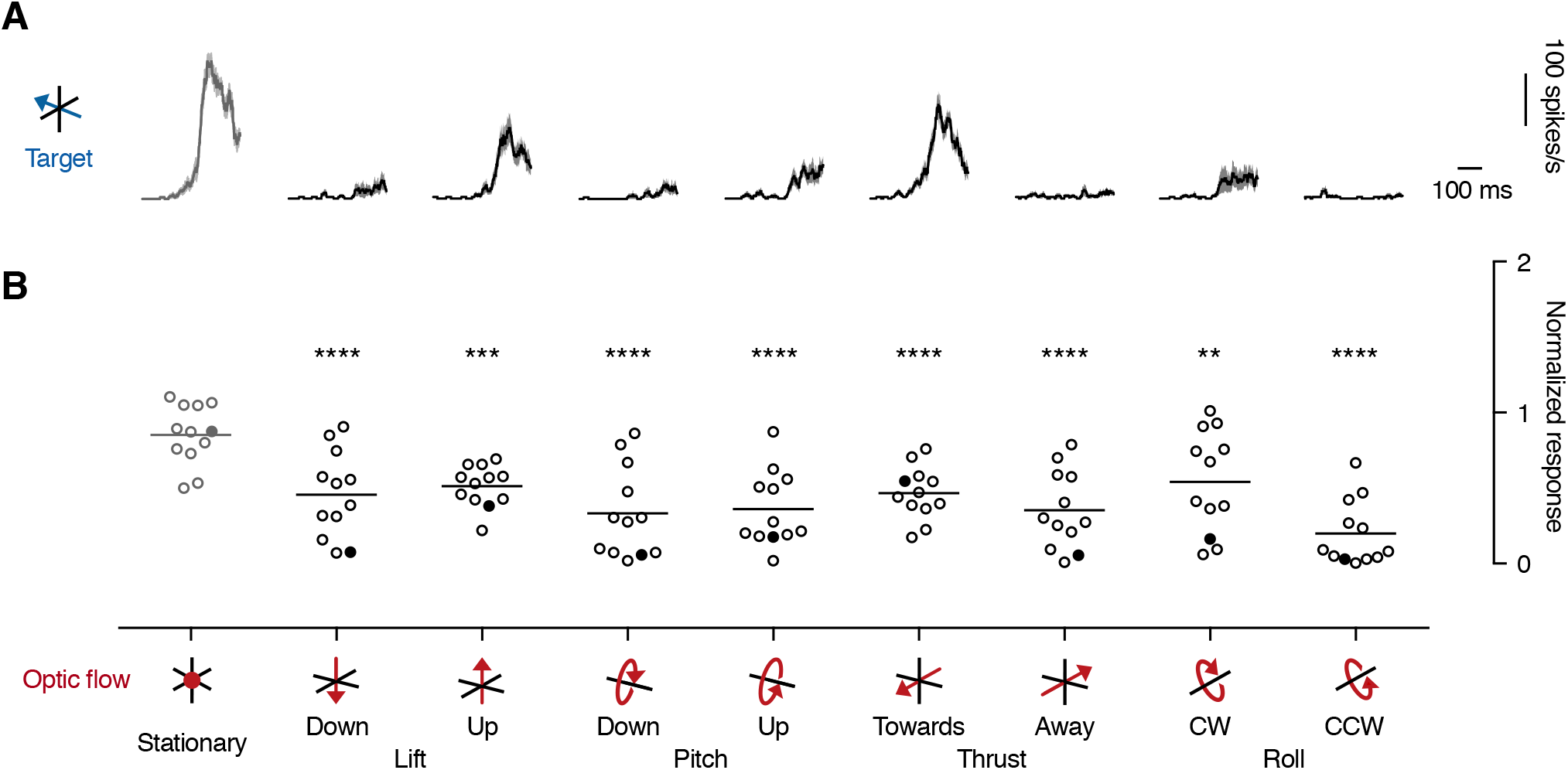
The TSDN response to target motion is attenuated by perpendicular optic flow. **A)** The response from one TSDN to a target traversing a stationary sparse dot field (left, grey, replotted from Fig. 2C), or different types of optic flow (red pictograms). All histograms shown as mean ± sem (n=18) at 1 ms resolution, after smoothing with a 20 ms square-wave filter. **B)** The mean response across TSDNs (N=12, same neurons as Fig. 2D) to a target traversing a stationary sparse dot field (left, grey, replotted from Fig. 2D), or different types of optic flow. Stars indicate significance, one-way ANOVA followed by Dunnett’s multiple comparisons test (**P<0.01, ***P<0.001 and ****P<0.0001), done together with the data shown in Fig. 2D. Translations were simulated at 50 cm/s and rotations at 50°/s. The lines show the mean and the filled data points correspond to the neuron in panel A.

We next looked at roll optic flow, which creates no local motion straight ahead of the fly, but opposite direction local motion in the dorsal and ventral visual fields. For example, during counter-clockwise roll (“Roll CCW”, Fig. 4) the target and the background optic flow move in the same direction through the dorsal receptive field, but in opposite directions in the ventral visual field. Therefore, the TSDN would receive a combination of maximum and minimum vector divergence signals from the “dorsal” and “ventral” parts (as shown in Fig. 3). We found that the response attenuation was stronger against counter-clockwise roll (75.3%, “Roll CCW”, Fig. 4), than against clockwise roll (31.9%, “Roll CW”, Fig. 4).

Thus, our data (Fig. 2, 4) support the notion that the level of vector divergence influences the TSDN response. To explore this in further detail, we recorded from TSDNs that respond robustly both to horizontal (grey, Fig. 5) and vertical target motion (black, Fig. 5). We found that when the target moved horizontally, the TSDN response was strongly attenuated against syn-directional sideslip (2^nd^ column, Fig. 5C), less attenuated against lift in either direction (last two columns, Fig. 5C), and strongly facilitated when displayed against counter-directional sideslip (3^rd^ column, Fig. 5C), consistent with previous results (Fig. 2, 4). In the same TSDN neurons, we next moved the target vertically. We found that the TSDN response was completely attenuated against syn-directional lift (last column, Fig. 5D), and less attenuated against sideslip in either direction (2^nd^ and 3^rd^ column, Fig. 5D), which moves perpendicular to the target. This supports the suggestion that maximum TSDN response attenuation is generated at minimum vector divergence between the target and background. However, when the vertical target was displayed against counter-directional lift (4^th^ column, Fig. 5D), there was no response facilitation. This suggests that maximum vector divergence on its own is not enough to explain the TSDN response facilitation to target motion.

**Figure 5.**
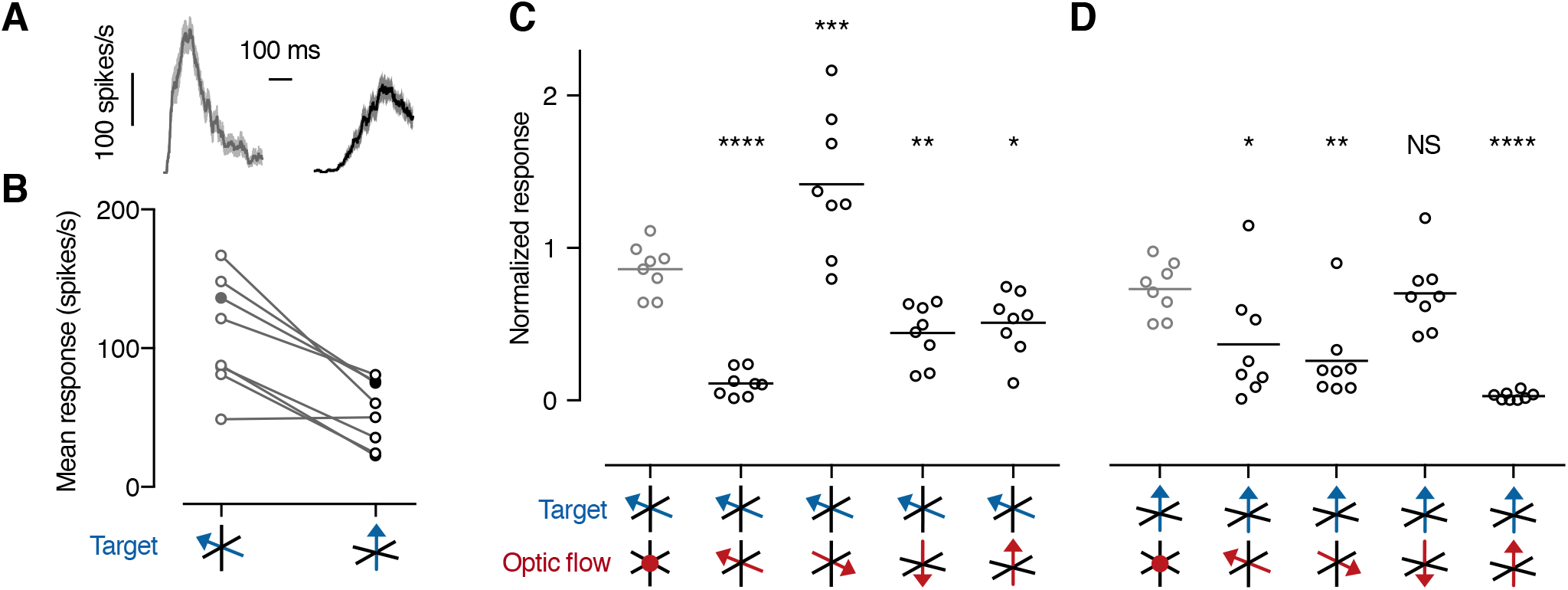
Vector divergence between optic flow and target. **A)** The TSDN response to a target traversing a white background horizontally (left, grey) or vertically (right, black). Histograms shown as mean ± sem (n=18) at 1 ms resolution, after smoothing with a 20 ms square-wave filter. **B)** The mean spiking response of 8 TSDNs to targets moving horizontally (grey, left) or vertically (black, right) over a white background. The filled data points correspond to the neuron in panel A. **C)** TSDN responses to targets moving horizontally across sideslip or lift, in the same direction as the target, or perpendicular to the target, as indicated by the pictograms. Significance shown using one-way ANOVA followed by Dunnett’s multiple comparisons test, with *P=0.0232, **P=0.006, ***P=0.0003, ****P<0.0001. **D)** TSDN responses to targets moving vertically across sideslip or lift, in the opposite direction to the target, or perpendicular to the target, as indicated by the pictograms. Significance shown using one-way ANOVA followed by Dunnett’s multiple comparisons test, with *P=0.0198, **P=0.0021, ****P<0.0001. Same N=8 in panels B-D.

### Responses to ON-OFF edges consistent with 1-point correlation

TSDNs have been proposed to get their input from STMDs (35). As opposed to our TSDN data, hoverfly STMD responses are rarely suppressed by syn-directional background motion, and not facilitated by counter-directional background motion (21). As there are many other target tuned neurons in the fly optic lobes (e.g. 33, 36–39) it is possible that TSDNs do not get their input from STMDs. We can investigate the potential input using the underlying target tuning mechanisms, which can be distilled down into three fundamentally different concepts. For example, visual neurons can become target tuned by receiving inhibitory feedback from the widefield system (39–41), or by using center-surround antagonism together with rapid adaptation (38, 42). Alternatively, they can use an elementary STMD model, which is tuned to the unique spatio-temporal profile of a moving target, with a dark contrast change (OFF) from the leading edge followed by a bright contrast change (ON) by the trailing edge. Importantly, while the first two mechanisms rely on comparisons from neighboring points in space, the elementary STMD compares input from one point on space (38, 43, 44). Therefore, the first two models will respond similarly to the motion of a target, to the motion of a leading OFF edge, and the motion of a trailing ON edge (black, Fig. 6, redrawn from (38, 45)). In contrast, the elementary STMD model only responds strongly to the target (grey, Fig. 6, redrawn from (45)).

**Figure 6.**
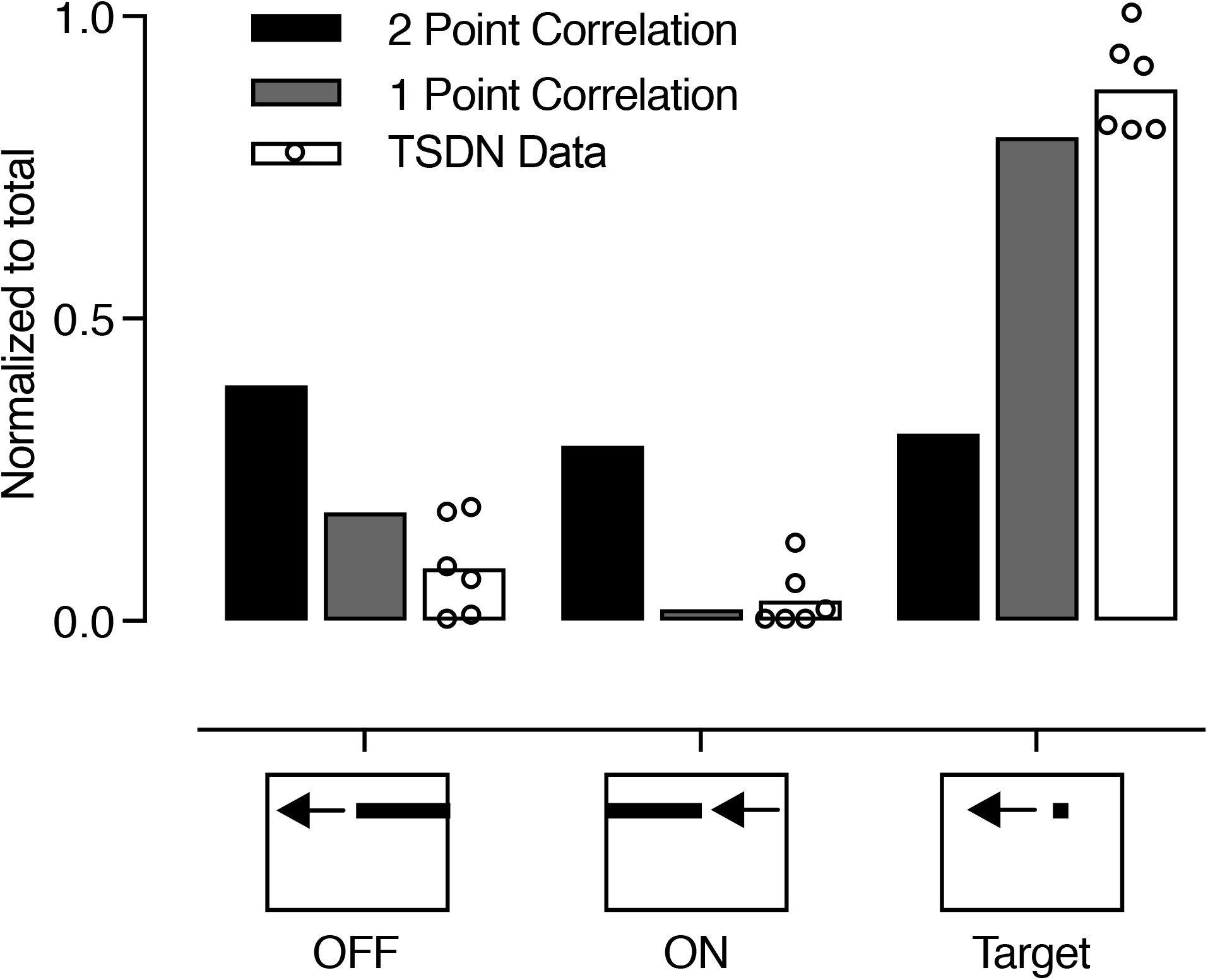
Elementary STMD input to TSDNs. Responses to a leading OFF edge, trailing ON edge, or a complete black target, with a side of 3°, traversing a white background at 130°/s. The black data show the predicted output from a motion detector that compares luminance changes from at least 2 points in space. Data replotted from (45), after normalizing to its own sum. The grey data show the predicted output from an elementary STMD (ESTMD), which compares luminance changes from one point in space. Data replotted from (45), after normalizing to its own sum. The white data show the TSDN response to the same three stimuli (N=6) after normalizing the data from each neuron to its own sum.

Our results show that TSDNs do not respond well to a leading OFF edge, or to a trailing ON edge (white, Fig. 6). However, a complete target, where the leading edge is rapidly followed by a trailing edge, gives a robust TSDN response (white, Fig. 6). Indeed, the physiological responses (white, Fig. 6) match the elementary STMD model output (grey, Fig. 6). Since STMD physiology also matches the elementary STMD model output (45, 46), this suggests that TSDNs receive input from STMDs.

## Discussion

We found that optic flow moving in the same direction as a target almost completely attenuated the TSDN target response (Fig. 2), even when the optic flow only covered a small part of the frontal visual field (Fig. 3). More strikingly, we found that optic flow in the opposite direction to target motion increased the TSDN response (Fig. 2), if the target moved horizontally (Fig. 5). We also found that perpendicular optic flow attenuated the TSDN target response (Fig. 3, 4), but less than syn-directional motion, suggesting that the vector divergence between the target and the background is important for response suppression. However, we found that vector divergence is not enough to explain the response facilitation (Fig. 3C).

We show for the first time, that TSDN responses to target motion are facilitated by counter-directional optic flow (Fig. 2). Such neural facilitation has not been seen in previous work in TSDNs (22) or STMDs (21, 26, 46), or other target tuned neurons in the optic lobes (33, 36, 39). Instead, in our previous work TSDN responses were significantly reduced when a background image with naturalistic statistics moved in the opposite direction to the target (22). Since the background used previously drive optic flow sensitive descending neurons as well as sinusoidal gratings do (22, 47), and as well as the sparse dot field used here (24), this suggests that facilitation is not generated by widefield motion in general. Indeed, this suggests that there is something fundamentally different about the optic flow used here, and the backgrounds in previous work. One difference is that we here use perspective distorted optic flow, where the individual features move faster the closer they are during translations, whereas during rotations they all move at the same angular velocity irrespective of distance from the viewer. However, we found that the facilitation was strong whether the target was displayed over sideslip or yaw (Fig. 2), suggesting that the relative motion of the features within the background optic flow is not important.

Another difference is that the background used here consisted of the coherent motion of thousands of “targets”, whereas previous TSDN work used a background image with naturalistic statistics (22). We currently do not know how STMDs, which are likely pre-synaptic to the TSDNs (Fig. 6, and see Ref. 35), respond to this stimulus. However, we do know that STMDs generate their ability to detect targets in clutter by being sharply tuned to the target’s unique spatio-temporal profile (43), with a dark-contrast OFF edge quickly followed by a bright-contrast ON edge (Fig. 6). Indeed, if naturalistic backgrounds contain small, high-contrast features, these often generate STMD responses (46). However, despite our optic flow consisting of “targets”, we saw no consistent TSDN responses even to preferred direction motion (e.g. “Sideslip Left”, “Yaw Left”, Fig. S1, Movie 5, 6). This suggests that TSDNs do not filter contrasting targets embedded within a background, like some STMDs do (21, 46).

It is thus difficult to determine whether the facilitation is driven by active facilitation onto the TSDN, or if it is inherited from upstream processes. If it is generated by neural input, and since the facilitation required frontal optic flow, but this did not have to spatially overlap with the TSDN receptive field or the target trajectory (Fig. 3A, C), the facilitation would come from a direction-selective neuron type with a frontal visual field. Since counter-directional lift did not facilitate the TSDN response if the target moved vertically rather than horizontally (Fig. 5C), this suggests that such a neuron would have to be most sensitive to horizontal motion, and less to vertical motion. Since some STMDs might respond to features within the optic flow used here (46), and furthermore, since there are some STMDs that respond both to target motion and to features within background motion (21), these could potentially play a role in this. Future investigation of STMD and TSDN responses are clearly required to explore the underlying mechanisms.

In contrast to the facilitation, we found that the sparse dot field stimulus suppressed the TSDN response to target motion when these moved in the same direction (Fig. 2). This response suppression could be driven by active inhibition from widefield motion detectors, as previously suggested (22), or inherited from upstream processes. We previously showed that TSDN responses to target motion are suppressed by background motion consisting of an image with naturalistic statistics, and that background stimuli that drive optic flow sensitive neurons suppress TSDN responses to target motion, whether spatially overlapping with the target trajectory or not (22). This suggests that TSDNs are suppressed by widefield motion in general. However, note that since we found maximum TSDN response suppression at minimum vector divergence, whereas perpendicular motion suppressed the response less (Fig. 4, 5; and see ref. 22), the response suppression has to contain some type of vector divergence comparison.

Nevertheless, our findings make behavioral sense. Prior to initiating target pursuit, male *Eristalis* hoverflies predict the flight course required to successfully intercept the target, based predominantly on the target’s angular velocity (48). To successfully execute an interception style flight course, the hoverfly turns in the direction that the target is moving (48). In doing so, the hoverfly creates self-generated optic flow in the opposite direction to the target’s motion. In this case the TSDNs would be facilitated (Fig. 2), which could be beneficial.

Once pursuit is initiated, if the pursuer drifts rightward due to e.g. a gust of wind, this induces leftward optic flow across the frontal retina, which would evoke a leftward optomotor response and/or gaze stabilizing turn. If the target is also moving leftward, in the direction of the corrective maneuver, this maneuver would also pursue the target and no TSDN signal would be required. Indeed, we found that the TSDNs were quiet under such conditions (“Left”, Fig. 2). However, if during the same pursuit the target instead moves rightward, opposite to the direction of the pursuer’s corrective maneuver, then the TSDNs would be strongly facilitated (“Right”, Fig. 2). Considering that TSDNs project to motor command centers in the thoracic ganglia (25, 31), this TSDN facilitation could then presumably override corrective turns. The TSDN response modulation by optic flow shown here could thus be beneficial for controlling behavioral output.

We only saw a response facilitation if the target moved horizontally, and not vertically (Fig. 5). This facilitation difference could be affected by the fact that the TSDNs that we recorded from responded better to horizontal motion than to vertical target motion (Fig. 5A). In addition, since we do not know where the TSDNs project to we do not know if the TSDNs control head movements, wing movements, or maybe both. Indeed, recent work suggests that gaze stabilizing turns by the head, and the wing optomotor response, are controlled independently, and by different visual components (12, 49, 50). Furthermore, the difference could be explained by different behavioral requirements to vertical and horizontal target deviations. For example, whereas horizontal deviations require the left and right wings to move differently, to induce a turn, vertical deviations would likely require the left and right wings to perform similar corrective behaviors. Finally, it is important to note that even if the responses to vertical target motion were not facilitated, neither were they suppressed (4^th^ column, Fig. 5D). There would thus still be a strong neural signal projected to the thoracic ganglia.

## Materials and Methods

### Electrophysiology

*Eristalis tenax* hoverflies were reared and maintained as previously described (51). For electrophysiology, a male hoverfly was immobilized ventral side up using a beeswax and resin mixture. A small hole was cut at the anterior end of the thorax to expose the cervical connective, which was then raised slightly and supported using a small wire hook, for insertion of a sharp polyimide-insulated tungsten microelectrode (2 MOhm, Microprobes, Gaithersburg, USA). The animal was grounded via a silver wire inserted into the ventral part of the hole.

Extracellular signals were amplified at 100x gain and filtered through a 10-to 3000-Hz bandwidth filter on a DAM50 differential amplifier (World Precision Instruments, Sarasota, USA), with 50 Hz noise removed with a HumBug (Quest Scientific, North Vancouver, Canada). The data were digitized via Powerlab 4/30 (ADInstruments, Sydney, Australia) and acquired at 40 kHz with LabChart 7 Pro software (ADInstruments). Single units were discriminated by amplitude and half-width using Spike Histogram software (ADInstruments).

### Visual stimuli

*Eristalis* males were placed ventral side up, centered and perpendicular to an Asus LCD screen (Asus, Taipai, Taiwan) at 6.5 cm distance. The screen had a refresh rate of 165 Hz, a linearized contrast with a mean illuminance of 200 Lux, and a spatial resolution of 2560 x 1440 pixels, giving a projected screen size of 155° x 138°. Visual stimuli were displayed using custom written software based on the Psychophysics toolbox (52, 53) in Matlab (Mathworks, Natick, USA).

TSDNs were identified as described (22, 24). In short, we mapped the receptive field of each neuron by scanning a target horizontally and vertically at 20 evenly spaced elevations and azimuths (24), to calculate the local motion sensitivity and local preferred direction. We then scanned targets of varying height through the small, dorso-frontal receptive fields (Fig. 3A) to confirm that each neuron was sharply size tuned with a peak response to targets subtending 3°-6°, with no response to larger bars, to looming or to widefield stimuli (22, 24).

Unless otherwise mentioned, targets were black and round with a diameter of 15 pixels, moving at a velocity of 900 pixels/s for 0.48 s. When converted to angular values and taking the small frontal receptive fields of TSDNs into account, this corresponds to an average diameter of 3° and a velocity of 130°/s (22). Unless otherwise stated, each target travelled in each neuron’s preferred horizontal direction (i.e. left or right) and across the center of its receptive field. Between repetitions, we varied the target elevation slightly, to minimize habituation (22). There was a minimum 4 s between stimulus presentations. Stimulus order was randomized.

Optic flow was generated as previously described (24). Briefly, the optic flow consisted of a simulated cube with 4 m sides, filled with 2 cm diameter spheres at a density of 100 per m^3^, with the hoverfly placed in the center. The coherent motion of these ca 6400 spheres around the hoverfly was used to simulate self-generated optic flow. The ca 1200 spheres anterior to the hoverfly were projected onto the screen, with their size indicating the distance from the hoverfly. Circles closer than 6 cm were not displayed. Six types of optic flow were simulated: three translations at 50 cm/s (sideslip, lift and thrust) and three rotations at 50°/s (yaw, pitch and roll). Unless otherwise stated optic flow was displayed for 0.48 s prior to the target. Both target motion and optic flow disappeared simultaneously.

In most experiments the optic flow covered the entire visual display. In some experiments, we limited the spatial extent of the optic flow, into 4 spatial positions (Fig. 3A). TSDN receptive fields tend to be located slightly offset from the visual midline, with preferred direction of motion away from the midline (Fig. 3A). We defined the lateral parts of the display as either ipsilateral or contralateral based on the preferred direction of each TSDN.

### Data analysis and statistics

We recorded from 30 TSDNs in 30 male hoverflies. We kept data from all TSDNs that showed a robust response to a target moving over a white background (Fig. 1, Movie 1, 2). We repeated this control throughout the recording, and only kept data from neurons that responded consistently. We only kept data from experiments with a minimum 9 repetitions. The data from repetitions within a neuron were averaged, and shown as spike histograms (mean ± sem) with 1 ms resolution, after smoothing with a 20 ms square-wave filter. For quantification across neurons, we calculated the mean spike rate for each neuron from the spike histogram for the duration of target motion, after excluding the first and last 40 ms of each 0.48 s target trajectory (dotted boxes, Fig. 1B, C), unless otherwise indicated. We normalized the responses to each neuron’s own mean response to a target moving over a white background.

Data analysis was performed in Matlab and statistical analysis in Prism 7.0c for Mac OS X (GraphPad Software, San Diego, USA). Throughout the paper *n* refers to the number of repetitions within one neuron, and *N* to the number of neurons. The sample size, type of test (paired t-tests or one-way ANOVAs, followed by Dunnett’s post hoc test for multiple comparisons) and P value is indicated in each figure legend. All data have been deposited to DataDryad.

## Acknowledgements

We thank current and past lab members for constructive feedback, and the Botanic Gardens of Adelaide for their ongoing support. Malin Thyselius provided the hoverfly pictogram in Fig. 2. Our research was funded by the US Air Force Office of Scientific Research (AFOSR, FA9550-19-1-0294), the Australian Research Council (ARC, DP170100008, DP180100144 and FT180100289), and the Flinders Foundation.

## Supplementary Information

**Figure S1. TSDNs do not respond to optic flow**

**A)** To determine the TSDN response to the optic flow we first quantified each neuron’s spontaneous rate, i.e. the spikes generated when viewing a white background, and used this as a threshold. For each neuron we then quantified the number of trials in which the response to the optic flow was above this threshold. This was converted to a percentage of responses. The data here show this percentage of responses across 12 TSDNs as mean ± sem. **B)** In each neuron, we quantified the mean response for those trials that were classified as responders (in panel A). This mean response was normalized to each neuron’s mean response to a target traversing a white background, as throughout the rest of the paper. Data are shown as mean ± sem (N=12).

***Movie 1. TSDN response to a target over a white background***

The movie depicts the stimulus as displayed on the screen, with the hoverfly positioned centered and perpendicular to the screen. The hoverfly was positioned ventral side up, but we have rotated the movie for display purposes so the dorsal side is up. The colored lines show the outline of the receptive field, mapped as described previously (24). The red data at the bottom of the movie show the response of an example TSDN.

***Movie 2. TSDN response to a target over a white background***

Same as Movie 1, but slowed down 10 times.

***Movie 3. TSDN response to a target over stationary optic flow***

The movie depicts the stimulus as displayed on the screen, with the colored lines showing the outline of the receptive field. The red data at the bottom of the movie show the response of an example TSDN. In this case the optic flow was stationary, and appeared 0.5 s before the target.

***Movie 4. TSDN response to a target over stationary optic flow***

Same as Movie 3, but slowed down 10 times.

***Movie 5. TSDN response to a target over syn-directional sideslip***

The movie depicts the response of the same example TSDN to the same target motion as in previous movies, but now traversing sideslip appearing 0.5 s before the target. The hoverfly was positioned at a distance of 6.5 cm. At this viewing distance, the resulting optic flow simulates the type of optic flow that would be generated by the hoverfly side-slipping through the world at 50 cm/s. The simulated optic flow consisted of ca 6400 spheres, of which roughly 1200 are projected onto the screen at any one time.

***Movie 6. TSDN response to a target over syn-directional sideslip***

Same as Movie 5, but slowed down 10 times. Note that many of the optic flow “targets” of the correct size move through the receptive field (colored contour lines).

***Movie 7. TSDN response to a target over opposite direction sideslip***

The movie depicts the response of the same example TSDN to the same target motion as in previous movies, but now traversing sideslip in the opposite direction to the target.

***Movie 8. TSDN response to a target over opposite direction sideslip***

Same as Movie 7, but slowed down 10 times.

